# PBMC Treatment Significantly Changes Gene Expression Regulation in Horses

**DOI:** 10.1101/519769

**Authors:** Victor C. Mason, Tosso Leeb, Vinzenz Gerber

## Abstract

Expression quantitative trait loci (eQTLs) are context dependent, and therefore change between tissues, cell types, and after cell treatment. In addition, SNP positions and RNAseq counts must be updated after assembly of new reference genome sequences. Therefore, we remapped eQTLs with Matrix eQTL using the previously generated and publicly available data from four contexts of peripheral blood mononuclear cells (PBMCs) from European Warmblood horses to the EquCab3.0 reference genome, and used a linear mixed model in R to identify eQTLs with significantly different gene expression regulation in treated PBMCs when compared to no treatment (baseline). We found no evidence that SNPs associated with significant changes in gene expression between MCK and a treatment in PBMCs caused strong opposing regulatory effects. We identified canonical pathways with a significant number of genes in PBMCs with altered gene expression regulation when treated with lipopolysaccharides (LPS) and hay-dust extract (HDE). Significant pathways included RhoA signaling in LPS, as well as histamine degradation, cholesterol biosynthesis, FcγRIIB signaling, and others in HDE. Our results support previous research indicating that pathways altered between baseline and treatment of PBMCs in horses with LPS or HDE affect inflammatory responses through RhoA, B-cell signaling, IL-4 and IFN-γ, and histamine.

## 1. Introduction

Peripheral blood mononuclear cells (PBMCs) are cells with a single round nucleus (T-cells, B-cells, and natural killer (NK) cells, etc.) that are readily isolated from whole blood [1]. The transcriptional responses of PBMCs in four different *in vitro* contexts has been measured with RNAseq data in horses: no treatment (MCK) to represent baseline RNA expression, lipopolysaccharides (LPS) to mimic an inflammatory response, recombinant cyathostomin antigen (RCA) to mimic response to parasitic antigens, and hay-dust extract (HDE) to mimic severe equine asthma (SEA) (formerly known as recurrent airway obstruction, RAO) exacerbation in susceptible horses [2–5]. These RNAseq data were previously used to generate the horse transcriptome, and discover differentially expressed genes between horses with and without SEA [2–4]. The equine 670k SNP array was previously used to discover SNPs associated with SEA, and both SNP and RNAseq data were used to discover expression quantitative trait loci in horses (eQTLs) for the EquCab2 horse genome assembly [6–8].

A region of the genome containing a variant that influences the number of expressed RNA molecules from a gene is an eQTL. eQTLs are reproducible when the same conditions are applied to the same cell type from the same species [9,10]. However eQTLs are also context-dependent, and therefore eQTLs change depending upon environmental conditions, length or type of cell treatment, or cell types analyzed [11,12]. Therefore, analysis of many tissues and cell-types under various environmental conditions is required to understand context-dependent changes in gene expression regulation in various species.

Historically, eQTL studies analyzed multiple treatments of the same cells separately, determined significance based upon an alpha cutoff value, and searched for overlap between the lists of significant eQTLs. These studies provide lists of eQTLs present or absent in each context, but do not explicitly model changes between eQTLs across contexts and can be misleading due to variation in statistical power between contexts. Therefore, many recent studies have jointly modeled different contexts to ameliorate the differences in statistical power, and identify eQTLs unique and shared between treatments. Here, we used an interaction term in a mixed model to describe how eQTLs change between baseline (MCK) and treatment of PBMCs (LPS, RCA, HDE) in European Warmblood horses. Additionally, we updated previously published eQTLs to the EquCab3.0 reference genome, and identified pathways significantly enriched for genes with altered gene expression regulation between MCK and each PBMC treatment (LPS, RCA, or HDE) with *Ingenuity Pathway Analysis* (IPA) [13].

## 2. Materials and Methods

### An outline of the computational workflow is shown in figure 1

#### 2.1 Sample Information

Samples used in this study were previously collected, isolated, treated, and extracted as described in earlier publications [2,6–8,14,15]. Horses were kept in “low dust” environments before sample collection so that the SEA affected horses were in partial or full remission of SEA [2]. Horses were kept in stables with daily access to pasture all over Switzerland [2]. SEA horses received no prior treatment for SEA and a clinical exam was performed to rule out other systemic or localized infections [2].

DNA was previously extracted from PBMCs during two different studies [6,7]. PBMCs were previously treated, RNA extracted, and RNA sequenced (RNAseq) by Pacholewska et al. [15]. Pacholewska et al. followed the density gradient centrifugation procedure from Hamza et al. to isolate PBMCs and followed the treatment of PBMCs and RNA extraction method from Lanz et al. [2,14,15]. European Warmblood horses were selected as the breed to study because of the two warmblood families (Fam1 and Fam2) with high incidences of SEA [16]. eQTL analyses used DNA and RNA from 82 European Warmblood horses (40 with SEA, and 42 healthy). Ages of SEA (mean = 16.7, min = 10, max = 24, units = years) and healthy controls (mean = 17.8, min = 6, max = 32, units = years) were comparable. These 82 horses belong to three familial cohorts, two half-sibling (half-sib) families with 17 individuals (Fam1) and 15 individuals (Fam2) respectively, and 50 unrelated horses. The sires of Fam1 and Fam2 both had SEA. Unrelated horses are not part of Fam1 or Fam2, and do not show strong patterns of population structure within the group (S3 Fig). Unrelated horses are minimally two generations removed from one another (unrelated at the grandparent level) [7].

#### 2.2 SNP coordinate conversion to EquCab3.0 and filtration

We used the NCBI remap API to convert a VCF file of imputed SNPs from EquCab2 to a VCF file with coordinates for EquCab3. The VCF file used is available on the European Variant Archive (EVA) as project accession: PRJEB23301. This file was split into separate files with 5,000 lines each with a python script. Then each file was submitted to the NCBI remap API with remap_api.pl to convert the SNP coordinates. We used the parameters --mode asm-asm and converted --from GCF_000002305.2 to --dest GCF_002863925.1, and specified –in_format vcf and –out_format vcf. Meta-data in the output VCF file was not properly preserved, and therefore we replaced the genotypes in the output file with the correct values from the input files with a python script.

SNPs with a minor allele frequency less than 0.05 were removed. We removed SNPs that deviated strongly from HWE p-value < 1e-6 when only including healthy individuals. SNPs were filtered with vcftools v0.1.14 [17]. PCA plots based upon SNP genotypes was previously published [8].

#### 2.3 RNA sequences remapped to EquCab3.0 and gene expression counts

RNA sequences were remapped to the EquCab3 genome with STAR v.2.5.3a [18]. We set the parameters for STAR as follows: --outFilterMultimapNmax 50 --seedSearchStartLmax 25 -- alignIntronMin 20 --alignIntronMax 1000000 --alignMatesGapMax 100000 --sjdbGTFfeatureExon exon --sjdbGTFtagExonParentTranscript Parent --sjdbGTFtagExonParentGene gene -- outFilterMismatchNmax 4 --outFilterType BySJout SortedByCoordinate --outSAMstrandField intronMotif.

The counting of and normalization of RNA molecules was done following Mason et al. (2018) with minor changes [8]. We defined gene features with the NCBI annotation (release 103) of the horse reference genome sequence EquCab3.0 (Assembly accession: GCF_002863925.1). We specified desired features to be all transcripts of genes with the transcriptsBy() function in the Bioconductor R library *GenomicFeatures* [19]. We counted the number of RNA reads that aligned to all transcripts of each gene with the summarizeOverlaps() function in the Bioconductor R *GenomicAlignments* library [19]. We simplified the count matrix to have one feature per gene, making genes (not transcripts of genes) the RNAseq count feature. In the summarizeOverlaps function we specified ‘mode = “Union”, singleEnd = FALSE, ignore.strand = TRUE, fragments = TRUE’. We counted each treatment separately, and required genes to have at least one RNAseq read aligned to the gene in one individual. We normalized read counts to make them comparable across individuals in *DESeq2*, and then exported them to calculate a mean read count cutoff with the KS test statistic [8]. Genes with mean normalized read counts below this mean count threshold removed from analysis for each treatment separately. After trimming the number of genes, the gene expression raw counts were again normalized and then variance stabilized with the varianceStabilizingTransformation() in DESeq2 once for each treatment separately [20]. PCA plots of the variance stabilized gene counts were generated with *DESeq2* (Fig S01). No individuals were identified to have aberrant expression profiles after analyzing the PCA plots, therefore no individuals were removed based upon expression profiles (Fig S01).

#### 2.4 eQTL analyses

We performed four multivariate linear models with *Matrix eQTL* (one for each context) to determine presence or absence of eQTLs in each treatment, and one mixed linear model to detect significant interaction terms representing significant changes in gene expression regulation between MCK and each treatment respectively.

##### 2.4.1 Multivariate linear models with *Matrix eQTL*

eQTLs for each treatment were detected with *Matrix eQTL* [21]. Local eQTL relationships within 500,000 bp upstream and 500,000 bp downstream of each gene‘s transcription start site were tested for 833,937 SNPs and 13,849 genes. eQTLs with FDR values less than 0.05 were considered significant. Significant eQTLs from these analyses were used to determine presence or absence of eQTLs for each treatment. The following multivariate model was used for the analyses.

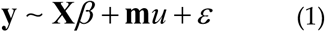

In equation two, **y** represents the dependent variable (normalized and variance stabilized gene expression), **X** is an incidence matrix for fixed effects intercept, age, sex, Fam1, Fam2, and disease status, *β* is the solution for the fixed effects intercept, age (in years), sex, Fam1, and Fam2, **m** is a vector of SNP marker genotypes, *u* is the SNP marker effect, and *ε* are the residuals.

##### 2.4.2 Mixed linear model analysis in R

The mixed model was ran on all local eQTL relationships within 500,000 bp upstream and 500,000 bp downstream of each gene‘s transcription start site were tested for 833,937 SNPs and 13,849 genes. We wrote a program in R that implemented a mixed model to jointly model one untreated baseline group, and three treatments of peripheral blood mononuclear cells (PBMCs) in European Warmblood horses to describe significant changes in eQTLs between baseline and the three treatments of PBMCs. The mixed model included an interaction term between genotype and context (treatement), corrected for effects due to population structure with binary variables Fam1 and Fam2, and we included a random intercept for each individual in R v.3.4.2 with R library hglm v.2.1-1 [22,23].

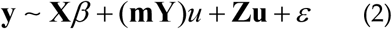

In equation two, **y** is the dependent variable representing the residuals of the trimmed, normalized, and variance stabilized gene expression counts. In equation one, **X** is an incidence matrix for fixed effects intercept, age, sex, Fam1, Fam2, disease status, SNP genotype (0, 1, or 2 for each individual: 0 is homozygous reference allele, 1 is heterozygous, and 2 is homozygous alternative allele), LPS context, RCA context, and HDE context, *β* is the solution for the fixed effects intercept, age (in years), sex, Fam1, Fam2, disease status, SNP genotype, LPS context, RCA context, and HDE context, **m** is a vector of SNP marker genotypes, **Y** is an incidence matrix that identified which gene expression values were present in which context (LPS, RCA, and HDE), *u* is the interaction effect of **m* Y**, **Z** is an incidence matrix which identified the repeated gene expression measurements (one from each context) for each individual, **u** is the vector of random individual effects, and *ε* is the random residuals.

#### 2.5. Removing associations with outlier individuals for the mixed model

Prior to multiple testing correction, we removed all models where with outlier individuals. Outlier individuals were identified by the R module hglm as “influential observations”. Influential observations were identified with the value “$bad” in the R model object of class hglm.

#### 2.5 Multiple testing correction for the mixed model

All raw p-values were corrected for multiple testing with EigenMT [24]. We ran EigenMT for each chromosome and for each covariate of interest. EigenMT also selects the best eSNP for each gene. We set the window size to include 200 SNPs, the variance threshold to 0.99, the *cis* distance to 500,000 (equivalent to 1Mb window), and considered results as significant if the adjusted p-value was < 0.05.

#### 2.6 Pathway analysis

A core analysis in Ingenuity Pathway Analysis (IPA) v.01-13 was run on all genes from eQTLs with significant interaction terms between baseline (MCK) and each of the three treatments of PBMCs (LPS, RCA, and HDE) [13]. Results were considered significant if the p-value was less than 1e-2. Results discussed are from the canonical pathway analysis. Gene names were mapped onto human, mouse and rat. We required the relationship between molecules to be direct and experimentally observed.

#### 2.7 Data availability

RNAseq data is deposited in the European Nucleotide Archive (ENA), and can be accessed at: http://www.ebi.ac.uk/ena/data/view/PRJEB7497 (project ID: PRJEB7497). Imputed SNP genotypes are submitted to European Variant Archive (EVA), project accession: PRJEB23301. Relevant python, R, and bash code is available on GitHub: https://github.com/VCMason.

## 3. Results

### 3.1 EquCab2.0 vs EquCab3.0

Gene expression counts in MCK were similar between EquCab2 and EquCab3 (r2 = 0.96) (Fig S02).

### 3.1 Changes in gene expression regulation by genotype due to PBMC treatment

An eQTL represents a significant association between a SNP‘s genotype and a gene‘s expression, while a significant interaction term represents a significant change in slope of an eQTL between MCK and each individual context. We detected 72,364 MCK, 100,382 LPS, 72,030 RCA, and 90,998 HDE significant eQTLs through *Matrix eQTL* analyses (Tables S1-S4). We detected a significantly different genotypic effect on gene expression between MCK and LPS, RCA, or HDE (a significant SNPxTreatment interaction term) for 1,057, 938, and 1,847 genes respectively (Tables S5-S7). To be kept for further analyses, we required each gene SNP pair with a significant interaction term (Tables S5-S7) to also be a significant eQTL in 1) MCK, 2) a treatment (LPS, RCA, or HDE), or 3) both MCK and treatment (Table 1, & S1-S4). This reduced the list of eQTLs analyzed to 144 for LPS, 60 for RCA, and 213 for HDE (Fig 2). Biologically, these represent 1) eQTLs present in MCK but not present in a treatment, 2) eQTLs present in a treatment but not present in MCK, 3) eQTLs present in both MCK and a treatment with the same direction of effect, or 4) eQTLs present in both MCK and a treatment with opposing directions of effect (Fig 2) [12]. We report the number of eQTLs with significant interaction terms present in each of the four scenarios (Table 1).

**Table 1.** The numbers of eQTLs with similar or opposite directions of effect (DOE) when MCK is compared to a treatment (LPS, RCA, or HDE).

|  | MCK Only | Treatment Only | MCK & Treatment | Not MCK or Treatment | Total |
| --- | --- | --- | --- | --- | --- |
| MCKxLPS: Same DOE | 39 | 45 | 24 | 102 | 210 |
| MCKxLPS: Opposite DOE | 19 | 17 | 0 | 811 | 847 |
| MCKxLPS: Total | 58 | 62 | 24 | 913 | 1057 |
| MCKxRCA: Same DOE | 24 | 10 | 11 | 79 | 124 |
| MCKxRCA: Opposite DOE | 13 | 2 | 0 | 799 | 814 |
| MCKxRCA: Total | 37 | 12 | 11 | 878 | 938 |
| MCKxHDE: Same DOE | 62 | 38 | 71 | 234 | 405 |
| MCKxHDE: Opposite DOE | 39 | 3 | 0 | 1400 | 1442 |
| MCKxHDE: Total | 101 | 41 | 71 | 1634 | 1847 |

We found no eQTLs with opposing directions of effect when eQTLs were significant in both MCK and a treatment (black data-points) (Fig 2, Table 1). All data points with opposing directions of effect in figure 1 have one non-significant gene/SNP association in either MCK or a treatment (orange or blue data-points). Therefore, we found no evidence that SNPs associated with significant changes in gene expression between MCK and a treatment caused strong opposing regulatory effects. Rather, the majority of eQTLs with a significant difference in gene expression regulation between contexts resulted in either a complete loss of an eQTL, gain of an eQTL, or the eQTL was modified but maintained the same direction of effect.

**Figure 1.**
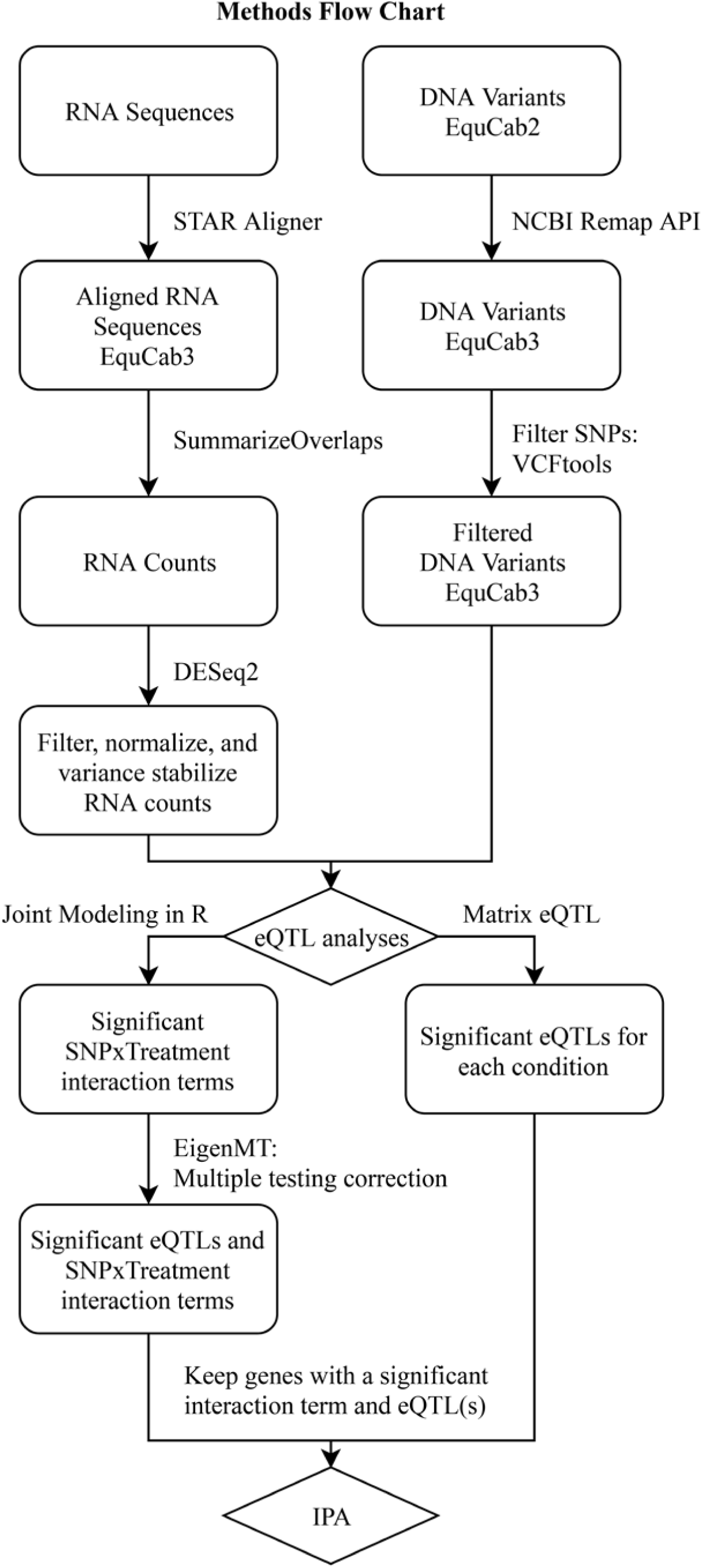

**Figure 2.**
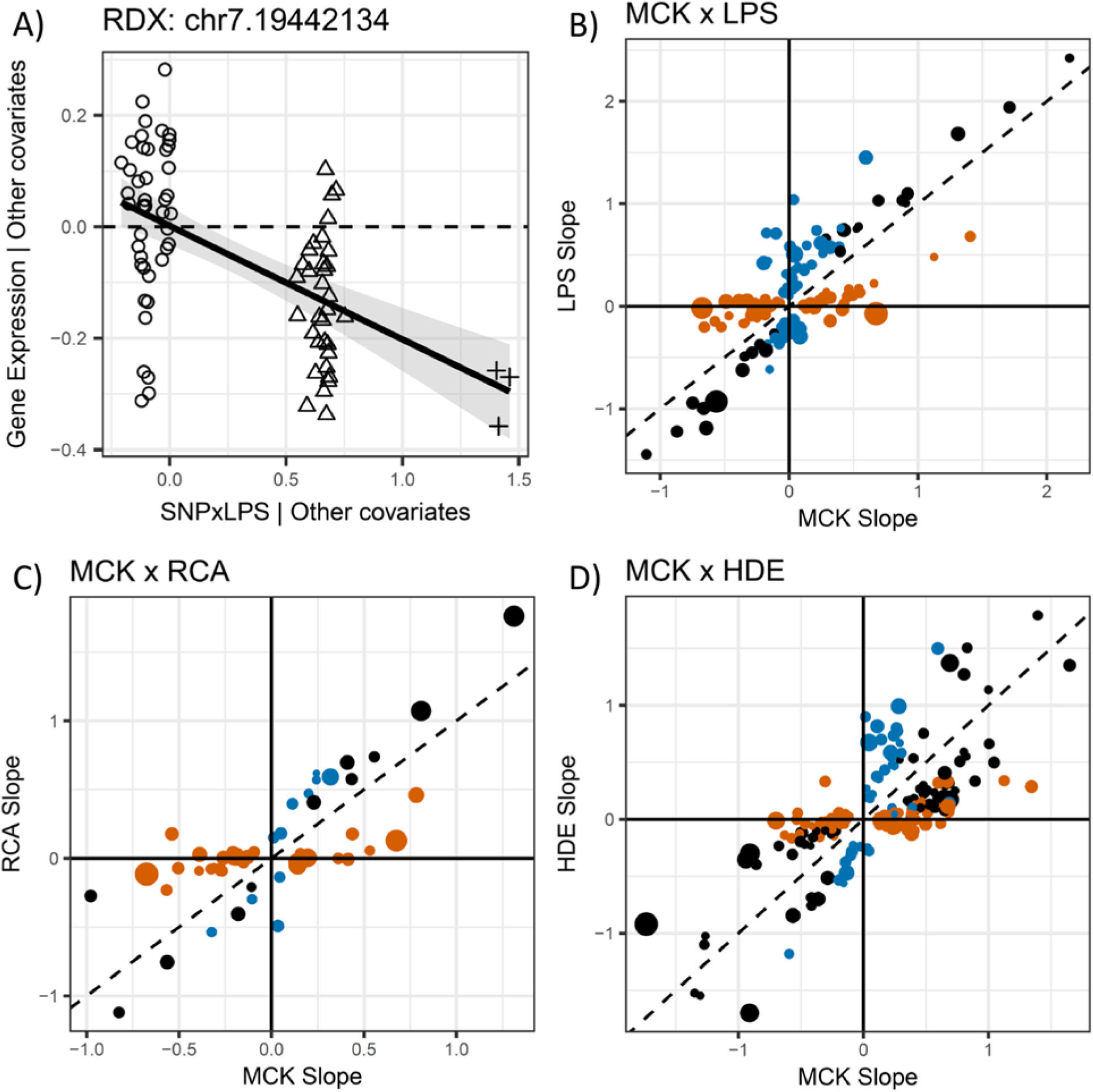

### 3.2 IPA

Three separate core analyses in IPA (one for each treatment) discovered biological pathways significantly (p-value < 0.01) enriched for genes with significantly (adjusted p-value < 0.05) altered gene expression regulation due PBMC treatment (Tables 2). Each treatment (LPS, RCA, and HDE) resulted in unique canonical pathways enriched for genes with altered gene expression regulation (relative to MCK) (Table 2).

**Table 2.**
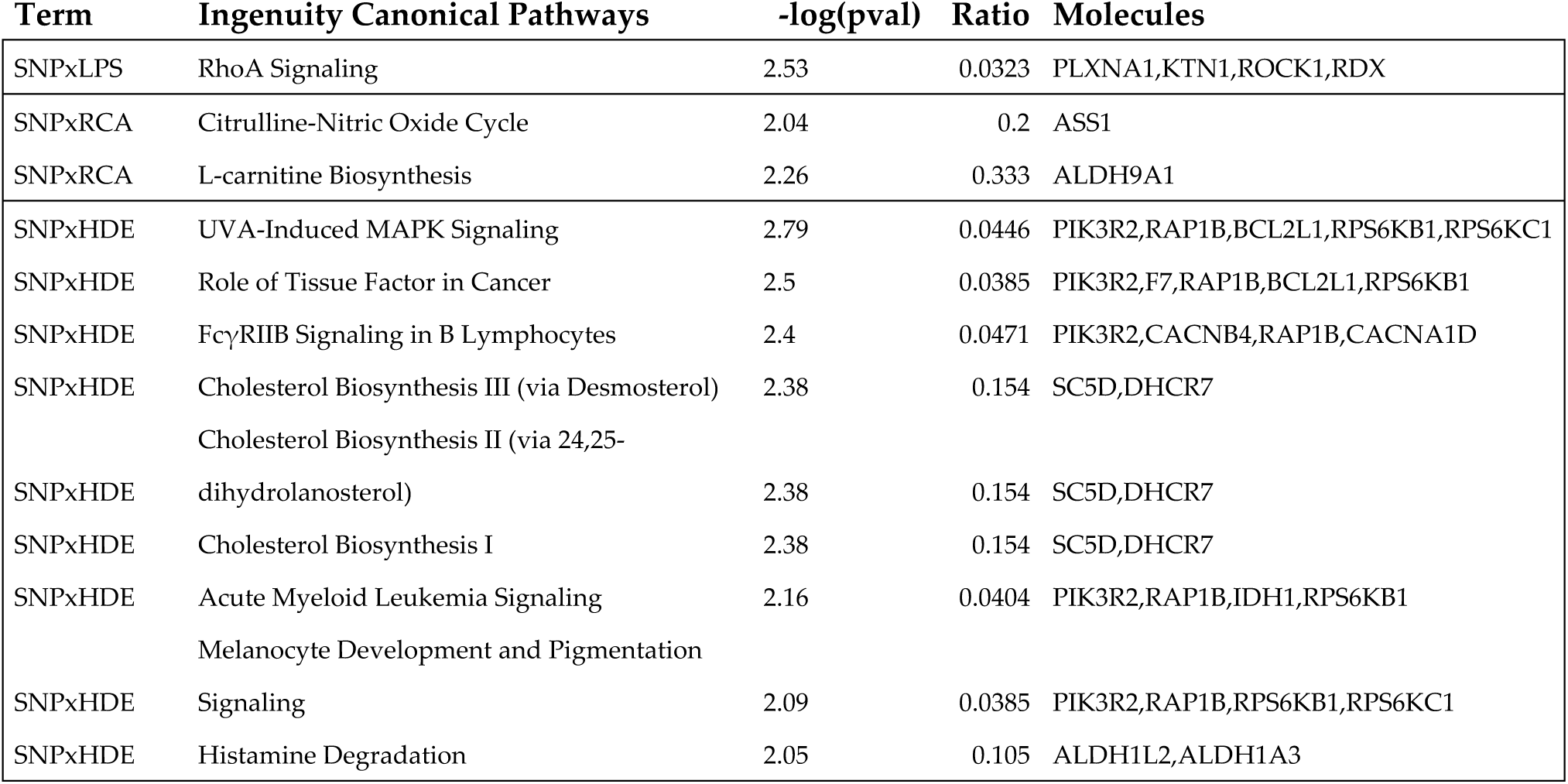
Significant canonical pathways with a significant number of genes with altered gene expression regulation.

## 4. Limitations

Horses with and without SEA were included in this study. We accounted for variation associated with disease status in the linear model, however some variation due to disease status (and other covariates) that is not explained by the fitted linear relationships may still be unaccounted for. Our method to determine presence or absence of eQTLs in each context might be improved with joint modeling of all treatments. We detected canonical pathways significantly enriched for genes with altered gene expression in PBMCs due to treatment. In the discussion we hypothesize about their relevance in horses, however we do not have further evidence about gene expression patterns in other tissues. Therefore, our hypotheses should be interpreted in an accordingly limited fashion.

## 5. Discussion

We discovered eQTLs associated with significant changes in gene expression regulation due to treatment of PBMCs in European Warmblood horses to identify gene pathways affected by treatment of LPS, RCA, or HDE. We removed variation in eQTLs correlated with age, sex, family structure, and disease (SEA) status to discover changes in eQTLs due to antigen treatment. By adjusting for these confounding factors, we focused our analysis on changes in gene expression regulation caused by antigen treatments. We expected LPS to influence genes involved in inflammatory regulation because it is present in the cell membrane of gram-negative bacteria, RCA to elicit an immune response as it is an antigen released from parasitic cyathostomins, and HDE to elicit allergic and inflammatory responses as HDE is derived from moldy hay. Therefore, we hypothesized that gene expression regulation altered by PBMC treatment would affect pathways involved in inflammatory and allergic immune responses. We detected significant canonical pathways for each treatment. However, the results for RCA may not be robust as only one gene was present in significant pathways. Therefore, we focused the discussion of significant canonical pathways on LPS and HDE treatments.

LPS altered gene expression for a significant number of genes in the RhoA signaling pathway. The RhoA signaling pathway is critical to pro-inflammatory responses and LPS/NF-κB signaling [25]. Depletion of RhoA in a human lung cancer cell line has been shown to significantly reduce the LPS-induced secretion of IL-6 and IL-8 [25]. Therefore, we hypothesize that RhoA also plays a role in the inflammatory response following PBMC stimulation with LPS in horses.

HDE significantly altered gene expression regulation in pathways involving cellular signaling, blood coagulation, vascular inflammation, and B cell signaling (Table 2). HDE influenced the histamine degradation pathway. Dietary histamines can cause allergic symptoms, gastrointestinal ailments, inflammatory responses, as well as a variety of other symptoms in histamine intolerant patients [26]. Histamine is present in respirable hay dust, and has been implicated as an irritating agent that contributes to respiratory problems, and is an inflammatory mediator secreted by IgE activated mast cells [27,28]. Cholesterol biosynthesis was also influenced by HDE treatment. Dietary cholesterol increases production of inflammatory indicators (IL-4 and IFN-γ) in lung lymphocytes in mice [29]. The two genes (*SC5D* and *DHCR7*) with altered gene expression regulation are required for the final steps of cholesterol biosynthesis (Table 2) [30,31]. Therefore, altered cholesterol metabolism could also contribute to the inflammatory symptoms in horses after inhalation of hay dust. FcγRIIB signaling in B lymphocytes was significantly affected by treatment of PBMCs with HDE. Fcγ receptors activate or inhibit inflammatory responses, and a proper balance is critical for normal immune response [32,33]. FcγRIIB is the only inhibitory Fcγ receptor, making it critical for controlled immune responses, and acts as a negative regulator of B cell activation.

## 6. Conclusions

We found no evidence that SNPs associated with significant changes in gene expression between MCK and a treatment in PBMCs caused strong opposing regulatory effects. Pathways influenced by PBMC treatment through changes in gene expression regulation include: RhoA signaling in LPS, as well as histamine degradation, cholesterol biosynthesis, FcγRIIB signaling, and others in HDE. Our results support previous research indicating that pathways altered between baseline and treatment of PBMCs in horses with LPS or HDE affect inflammatory responses through RhoA, B-cell signaling, IL-4 and IFN-γ, and histamine.

## Author Contributions

Conceptualization, VCM.; Methodology, VCM.; Software, VCM.; Validation, VCM..; Formal Analysis, VCM.; Investigation, VCM.; Resources, VG. and TL..; Data Curation, VCM.; Writing-Original Draft Preparation, VCM.; Writing-Review & Editing, VCM., VG., and TL.; Visualization, VCM.; Supervision, VCM., VG., And TL.; Project Administration, VCM., VG., and TL.; Funding Acquisition, VG.

## Acknowledgments

Thank you to Matthias Kraft and Dr. Vidhya Jagannathan for many stimulating conversations and helpful suggestions.

## Appendix A

**Figure 1.** Flow chart depicting informatic methods. **Figure 2.** Differences in eQTLs between baseline (MCK) and the three treatments of PBMCs B) LPS, C) RCA, and D) HDE. A) A partial regression plot for the interaction term SNPxLPS shows the difference in slope of an eQTL for gene RDX and the SNP located on chromosome 7 position 19442134 between MCK and LPS. The partial regression plot scales the eQTL in MCK to have a slope of zero and is represented by the dashed line. The eQTL in LPS is represented by the solid black line and the standard error of the eQTL is represented by the grey shading. Circles represent homozygous reference, triangles are heterozygous, and plus symbols represent homozygous alternative genotypes. The y-axis is the residuals of gene expression that is not explained by any covariates (excluding SNPxLPS). The x-axis is the residuals of product of SNP x LPS that is not explained by any other covariates. SNP is represented by values 0, 1, and 2 while LPS is 0 or 1. The three remaining figure sections represent the significance and change of gene expression regulation between baseline (MCK) and the three treatments of PBMCs B) LPS, C) RCA, and D) HDE. The area of the plot points are proportional to the –log10 of the p-values of the SNPxTreatment interaction term. The slopes of the eQTLs in MCK are plotted on the x-axis in B), C), and D). The slopes of the same eQTLs after treatment are plotted on the y-axis for B) LPS, C) RCA, and D) HDE. eQTLs significant in MCK, but not in a treatment B) LPS, C) RCA, or D) HDE are colored orange. eQTLs not significant in MCK but significant in a treatment B) LPS, C) RCA, or D) HDE are colored blue. eQTLs significant in both contexts B) MCK and LPS, C) MCK and RCA, and D) MCK and HDE are colored black. **Table S1.** Significant eQTLs (FDR < 0.05) in MCK calculated for the EquCab3 genome with *Matrix eQTL*. **Table S2.** Significant eQTLs (FDR < 0.05) in LPS calculated for the EquCab3 genome with *Matrix eQTL*. **Table S3.** Significant eQTLs (FDR < 0.05) in RCA calculated for the EquCab3 genome with *Matrix eQTL*. **Table S4.** Significant eQTLs (FDR < 0.05) in HDE calculated for the EquCab3 genome with *Matrix eQTL*. **Table S5.** EigenMT results reporting significant (Bonferroni correction < 0.05) SNPxLPS interaction terms. Table header labels are described as follows: ‘SNP’ is the chromosome.position of the SNP in the genome, ‘gene’ is the gene name, ‘beta’ is the slope of the interaction term, ‘t-stat’ is the t-statistic value, ‘p-value’ is the unadjusted p-value, ‘BF’ is the Bonferroni corrected p-values of the interaction term, ‘TESTS’ equals the effective number of SNPs used for the Bonferroni correction. **Table S6.** EigenMT results reporting significant (Bonferroni correction < 0.05) SNPxRCA interaction terms. Table header labels are described as follows: ‘SNP’ is the chromosome.position of the SNP in the genome, ‘gene’ is the gene name, ‘beta’ is the slope of the interaction term, ‘t-stat’ is the t-statistic value, ‘p-value’ is the unadjusted p-value, ‘BF’ is the Bonferroni corrected p-values of the interaction term, ‘TESTS’ equals the effective number of SNPs used for the Bonferroni correction. **Table S7.** EigenMT results reporting significant (Bonferroni correction < 0.05) SNPxHDE interaction terms. Table header labels are described as follows: ‘SNP’ is the chromosome.position of the SNP in the genome, ‘gene’ is the gene name, ‘beta’ is the slope of the interaction term, ‘t-stat’ is the t-statistic value, ‘p-value’ is the unadjusted p-value, ‘BF’ is the Bonferroni corrected p-values of the interaction term, ‘TESTS’ equals the effective number of SNPs used for the Bonferroni correction.

