## Supplementary figures and images for "PBMC Treatment Significantly Changes Gene Expression Regulation in Horses"

### Fig S01

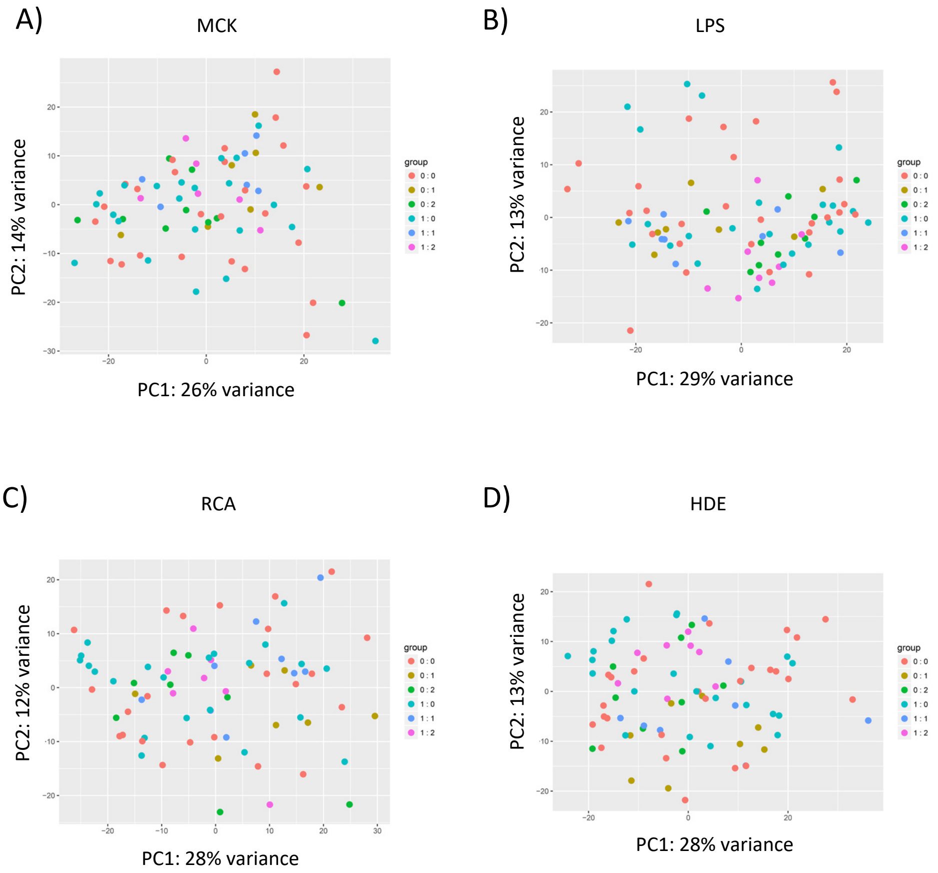

### Fig S02

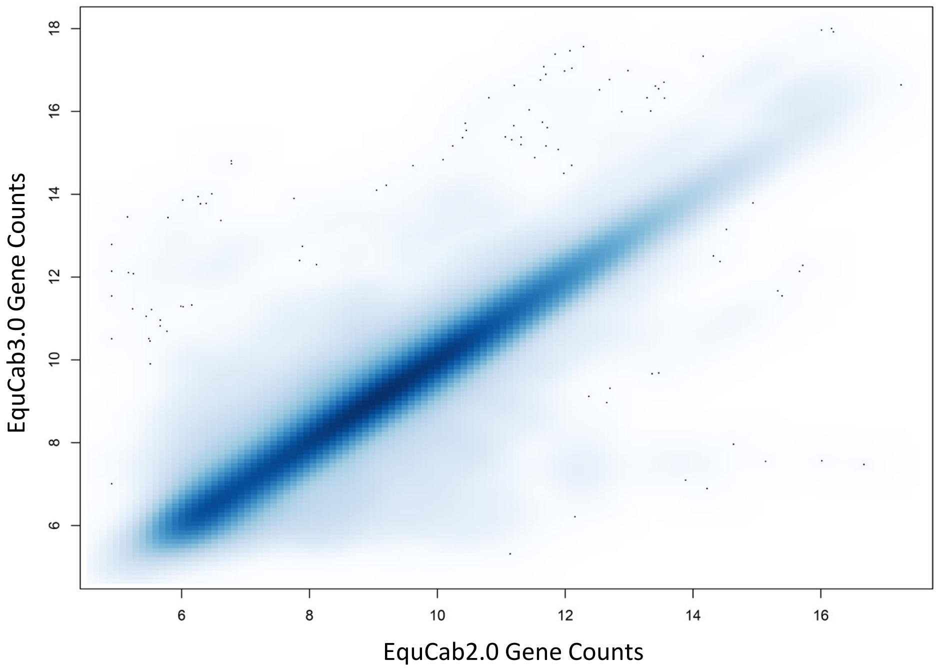
